# Improved assembly of noisy long reads by *k-*mer validation

**DOI:** 10.1101/053256

**Authors:** Antonio Bernardo Carvalho, Eduardo G. Dupim, Gabriel Nassar

## Abstract

Genome assembly depends critically on read length. Two recent technologies, PacBio and Oxford Nanopore, produce read lengths above 20 kb, which yield genome assemblies that are vastly superior to those based on Sanger or short-reads. However, the very high error rates of both technologies (around 15%-20%) makes assembly computationally expensive and imprecise at repeats longer than the read length. Here we show that the efficiency and quality of the assembly of these noisy reads can be significantly improved at a minimal cost, by leveraging on the low error rate and low cost of Illumina short reads. Namely, *k-*mers from the PacBio raw reads that are not present in the Illumina reads (which account for ~95% of the distinct *k-*mers) are deemed as sequencing errors and ignored at the seed alignment step. By focusing on ~5% of the *k-*mers which are error-free, read overlap sensitivity is dramatically increased. Equally important, the validation procedure can be extended to exclude repetitive *k-*mers, which avoids read miscorrection at repeats and further improve the resulting assemblies. We tested the *k-*mer validation procedure in one long-read technology (PacBio) and one assembler (MHAP/ Celera Assembler), but is likely to yield analogous improvements with alternative long-read technologies and overlappers, such as Oxford Nanopore and BLASR/DAligner.

> *“Thm: Perfect assembly possible iff*
>
> a. *errors random*
> b. *sampling is Poisson*
> c. *reads long enough 2 solve repeats.”*
>
> — Myers, 2014

> *“One chromosome, one contig.”*
>
> — Koren et al., 2012

## INTRODUCTION

Genome assembly quality depends on sequencing coverage, read accuracy, and read length (Nagarajan and Pop 2013; Myers 2016). Nowadays the cost per sequenced base is small, so in many cases coverage is no longer a major limiting factor, 100-fold coverage being routine in many projects. Such high coverages also reduce the importance of read accuracy, since errors can be removed by consensus while building contigs from the reads. Read length remains a critical factor. Its importance stems from repeated sequences, which in many cases cannot be properly assembled unless they are shorter than the read. For example, two identical copies of a 7 kb retrotransposable element would require reads bigger than their length for being fully assembled; shorter reads would produce a fragmented assembly. This limitation can be circumnavigated, but only partially, by mate-pair reads and other methods (Weber and Myers 1997; Nagarajan and Pop 2013; McCoy et al. 2014).

Sanger sequencing, the first practical technology for large-scale projects, produce reads between 500 bp to1 kb, which are accurate (error ratê1 %), but expensive, the price tag for a *Drosophila*-like genome being in the million dollars range (Drosophila_Community_Resources_Committee 2001). Second generation sequencing technologies (“SGS”) such as Illumina produce reads that are inexpensive, accurate (error rate ~1 %), but short (< 500 bp). Their low cost (*Drosophila* genome price tag: ~U$ 4000) allowed for an explosion of genome projects. However, due to their short read length, they produce very fragmented assemblies. Both Sanger and SGS require a huge investment of money and time if a “finished” genome is the target.

Two recently developed or improved technologies, PacBio and Oxford Nanopore, produce read lengths above 20 kb, which can yield genome assemblies that are vastly superior in contiguity to those based on Sanger or short-reads (Goodwin et al. 2015; Koren and Phillippy 2015; Loman et al. 2015). However, reads produced by both technologies have very high error rates (PacBio: ~15%; Oxford Nanopore: ~20%), and cannot be directly handled by current genome assemblers (see (Li 2016) for an exception). Instead, a “hierarchical assembly process” is used: first the raw reads are error-corrected by aligning them either to Illumina reads (“hybrid assembly”; (Koren et al. 2012), or among themselves (“self-correction”; (Chin et al. 2013)), and implementing some sort of consensus algorithm, which reduces the error rate to below 5%. The corrected reads are then assembled by normal OLC assemblers (*i.e*., designed for Sanger reads). Self-correction produces better assemblies (Koren et al. 2013) and is the current state-of-art but is computationally intensive, because the all-by-all alignment must be carried with rather high sensibility and specificity in order to detect real overlaps among the noisy reads. In practice bacterial genomes are easily assembled, but large genomes such as mammals still have nearly prohibitive computational costs (*e.g*., more than 800,000 CPU hours; (Berlin et al. 2015). It is also unclear how far can we go inside highly repeated regions such as heterochromatin (*e.g*., telomeres and centromeres), segmental duplications and tandem gene arrays (*e.g*., histone and rDNA clusters).

A very efficient approach to analyze DNA sequences is to decompose them into overlapping stretches of a fixed length of *k* bases, called *k-*mers. For example, overlapping reads can be detected because they share *k-*mers above a certain cut-off (Berlin et al. 2015). *k-*mers provide interesting insights into the above mentioned relationship between read accuracy and computational cost, as can be seen in the following real example (throughout this manuscript we set *k*=16, which is a typical value). The genome of the bacterium *E. coli* strain K-12 MG1655 has been fully sequenced and finished to high quality years ago, using Sanger reads (Blattner et al. 1997). More recently, it has been sequenced using Illumina and PacBio technologies at high coverage (77x and 94x respectively; (Kim et al. 2014); https://basespace.illumina.com). The genome itself has 4.64 Mbp, and hence contains approximately 4.64 million distinct *k-*mers, the vast majority of them occurring only once (bacterial genomes have few repetitive regions). The PacBio reads contain a **total** of 436 million *k-*mers (4.64 million *k-*mers times 94-fold coverage); if there were no sequencing errors, these *k-*mers would correspond to 4.64 million **distinct** *k-*mers, each one occurring on average 94 times. However, these reads actually contain 292,687,635 distinct *k-*mers (~293 millions); among these, 4,513,248 (1.5%) are correct (*i.e*., present in the finished *E. coli* genome), and the remaining 288,174,387 are sequencing errors (“error *k-*mers”; see Methods). As expected, the correct *k-*mers show up repeatedly, and their proportion among the **total** *k-*mers is 16.6%. On the other hand, most error *k-*mers are unique, because the chance that random errors create twice the same 16-mer sequence (or a pre-existing 16-mer) is small. Fig. 1 shows a graph of the *k-*mer frequency spectrum of the PacBio reads and also, for comparison, of Illumina reads. It is easier to consider first the Illumina reads (Fig. 1, left panels): the huge peak on the left contains rare *k-*mers that mostly resulted from sequencing errors; the next peak, located approximately at the sequencing coverage, corresponds to single-copy sequences in the genome; finally, smaller peaks on the right correspond to repetitive DNA (they are much more pronounced in repeat-rich genomes such as *Drosophila* and mammals). A similar pattern occurs with PacBio reads (Fig. 1, right panels), except that the error peak is much higher (note the Y-axis scale), and that the single-copy peak is strongly shifted towards the left (because so many *k-*mers were “lost” due to sequencing errors). Roughly similar values were obtained for other genomes; more typically, ~5% of the distinct *k-*mers of the PacBio reads are correct (Supplemental Table S1).

**Figure 1.**
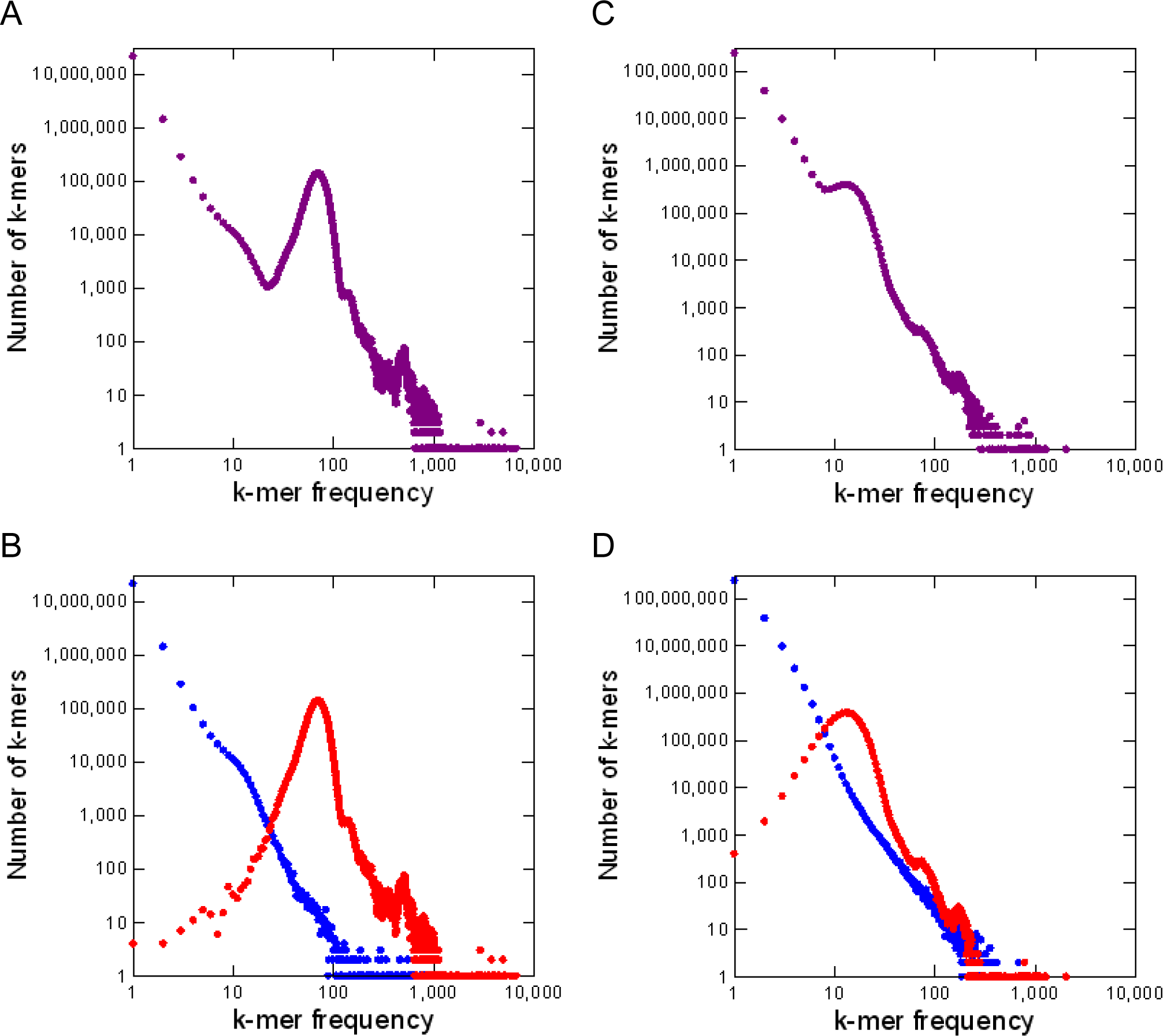
*k-*mer frequency distributions for Illumina and PacBio *E. coli* reads. Panel A, Illumina, all *k-*mers (*k*=16 in all panels); panel B, Illumina, with correct *k-*mers shown in red and error *k-*mers in blue. Note that most error *k-*mers have very low frequency. The peak at *k-*mer frequency ~ 70 corresponds to genomic single copy *k-*mers. Panels C and D, PacBio reads. Note the huge number of error *k-*mers. The reference list of valid *k-*mers came from the finished genome (see Methods).

The relevance of the data shown in Fig. 1 became apparent when one considers that all genome assembly algorithms are based on *k-*mer decomposition and comparison, and that at least at some steps they must track all distinct *k-*mers. As on average only ~5% of the distinct *k-*mers of the PacBio reads are correct (Supplemental Table S1), at some steps 95% of the computational resources such as memory and CPU time are wasted with *k-*mers that cannot indicate real read overlaps because they contain at least one wrong basis. The three main aligners for PacBio reads deal with this problem somewhat differently. BLASR, the first developed, originally aimed to align PacBio reads to a reference genome, but can also do the all-by-all alignment. It uses all *k-*mers and employs successive refinements of the alignment in order to detect true overlaps (Chaisson and Tesler 2012). The main limitation of BLASR is its low speed: it works well for bacteria and yeast genomes (4-12 Mbp), but is unpractical for genomes such as *Drosophila* (180 Mbp; it used 610,000 CPU hours in the all-by-all step; (Berlin et al. 2015)). DALIGNER employs a highly optimized code to perform similar tasks (Myers 2014). It is computationally intensive, and in practice requires large clusters to assemble *Drosophila*-like genomes. Its code in under active development; in a recent study DALIGNER was unstable with large genomes (Berlin et al. 2015). The third aligner, MHAP, reduces memory usage and computational time by sampling a random subset of *k-*mers (“sketch”) to detect candidate overlaps; larger sketch size results in more sensitivity, and in a higher computational cost. Typically sketch sizes range from ~500 to 1200 *k-*mers, with resulting sensitivities in the range of 60%-90% (Berlin et al. 2015). MHAP is the default aligner of the PBcR pipeline, which, after correcting the reads, feed them into the Celera Assembler. Currently PBcR is the pipeline that requires the smaller computational infra-structure: microbial genomes can be assembled in a 8-core desktop in a few hours or less, *Drosophila*-sized genomes are assembled in small servers (*e.g*., 24 cores, 64 Gb of RAM) in a couple of days, and mammalian size genomes require a large cluster.

Whatever the details of the overlapper algorithm, they all have to cope with a “needle-in-haystack” problem (*i.e*., to find true overlaps amidst a lot of sequencing noise), and in principle would work much better if the large number of “error *k-*mers” of the long, noisy reads could be identified at the outset, and ignored. We propose a simple solution to achieve this. Note that particularly in the case of Illumina reads the large peak on the left contain nearly no correct *k-*mer (Fig. 1); nearly all of them correct *k-*kmers are located to its right. This suggest an interesting possibility: in the absence of a finished genome, an accurate list of “valid *k-*mers” can be obtained from the Illumina reads by taking those *k-*mers that occur at least, say, 10 times (single-copy *k-*mers are expected to occur ~70 times on this dataset). In the *E. coli* example, if we use the Illumina-based list to validate the *k-*mers of the PacBio reads, we would miss only 145 correct *k-*mers (out of 4,513,248), and would incorrectly validate 0.1 % of the error *k-*mers (32,456 *k-*mers out of 288,174,387). Such Illumina “valid *k-*mer list” is inexpensive to produce, and may improve long-read assemblies by identifying in the long reads the *k-*mers that should be ignored. We tested this idea with PacBio reads, assembled by MHAP / Celera Assembler. Indeed, ignoring the *k-*mers which are outside the valid *k-*mer list resulted in a much higher sensitivity in overlap detection at a smaller computational cost, improved the read error correction and, more importantly, yield more contiguous assemblies of repeat-rich genomes and regions. Given that it addresses a general problem of the noisy long reads, this procedure is likely to improve assemblies produced by other technologies (Oxford Nanopore) and aligners (such as BLASR and DALIGNER).

## RESULTS

We implemented the *k-*mer validation in the MHAP overlapper as detailed in Methods, and tested its performance first in overlap detection, then in read error correction and finally on genome assembly. We used five model organisms; in four of them PacBio and Illumina reads from the same strain are available: a bacteria (*E. coli* strain K-12 MG1655; genome size of 4.64 Mbp), yeast (Saccharomyces cerevisae strain W303; 12.1 Mbp), worm (*Caenorhabditis elegans* strain Bristol N2; 103 Mbp) and flies (*Drosophila melanogaster* strain ISO1; ~180 Mbp). We also included the plant *Arabidopsis thaliana* (strain Ler-0; 135 Mbp), although in this case most of the Illumina reads came from different strains (mostly Ler-1), and were shallower (Supplemental Table S2; Supplemental Table S8). Finally, as a proof of principle, we applied the *k-*mer validation to a difficult region of the human genome. We operationally defined as valid *k-*mers all those with a frequency bigger than one seventh of the single-copy peak from Illumina reads (Supplemental Fig. S1 and Supplemental Table S2). This cut-off was chosen after a limited exploration (Supplemental Results).

### *k-*mer validation increases the sensitivity and the specificity of overlap detection

The MHAP program compares pairs of uncorrected PacBio reads, aiming to detect real overlaps while keeping false positives at a minimum. We compared the performance of the modified MHAP against the standard version (1.5b1) following the procedures of the original publication (Berlin et al. 2015). Namely, artificial PacBio reads were generated by applying the typical PacBio error rates (insertion: 10%; deletion: 2%; substitutions; 1%) to 10 kb segments of known genomes (we tested *E. coli*, yeast, *C. elegans*, and *Drosophila*, and also random DNA sequences). These segments were arranged as pairs with a 2 kb overlap; members of different pairs do not have any real overlap, but may contain similar sequences due to repetitive DNA. We measured sensitivity as the proportion of true overlaps that were detected (*i.e*., among members of the same pair). Overlaps between members of different pairs estimate the false positive rate (*i.e*., the specificity); this is more reliably done with random DNA sequences, because biological sequences almost always contain repeats that will inflate the false positive rate. We varied sketch size (the “num-hashes” parameter; (Berlin et al. 2015)) between 64 and 2048; this parameter is very important because it controls the trade-off of computational cost (CPU time plus memory usage) vs. sensitivity. All other parameters were kept fixed at their default values (*k-*mer size=16; num-min-matches=3; threshold =0.04). As shown in Fig. 2, *k-*mer validation caused a huge increase in sensitivity with *E. coli* data: at a sketch size of 512 (a typical value; (Berlin et al. 2015)), the standard MHAP detected 24% of the true overlaps (61 out 250), whereas with *k-*mer validation we got 95% (238 out 250). Other genomes and also random DNA sequences produce similar results (Supplemental Fig. S2). Finally, the improvement using the Illumina-derived list of valid *k-*mers is very similar to the one using the true *k-*mer list derived from the finished genome (Supplemental Fig. S3), which suggests that the former is an excellent proxy for the later.

**Figure 2.**
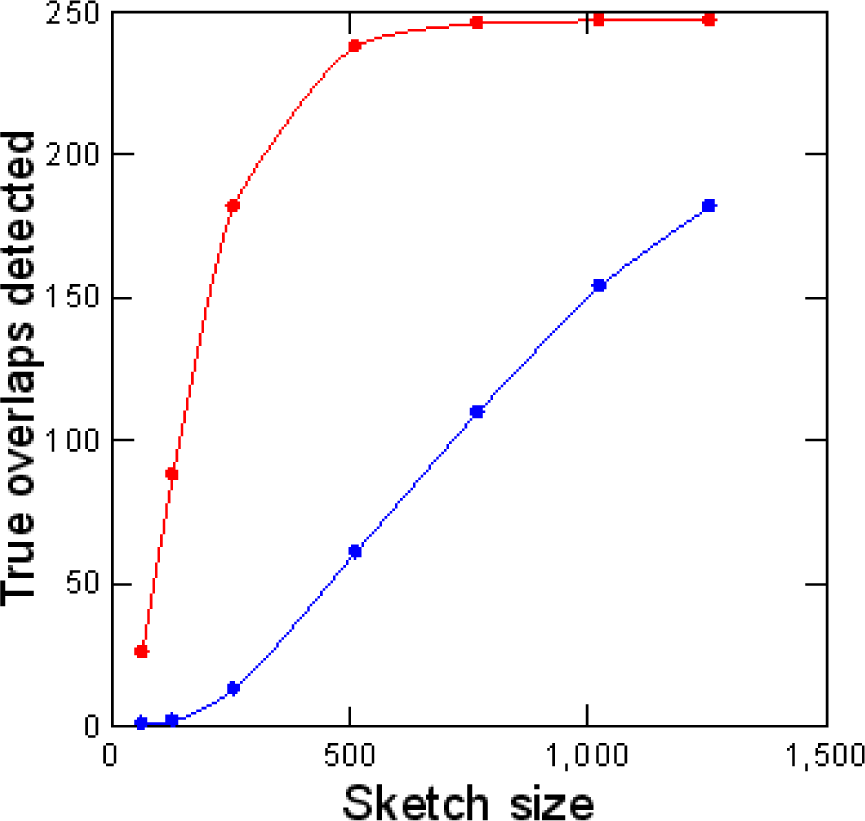
Sensitivity of read overlap detection with and without *k*-mer validation. Simulated PacBio reads from *E. coli* (250 pairs of 10 kb sequences with 2kb overlaps) were subjected to standard MHAP (blue) or MHAP with masking of low-frequency *k-*mers (red) for overlap detection. The reference list of valid *k-*mers came from Illumina reads.

It is interesting also to look at the false-positive rate, which estimates the specificity. The observation that false-positives are absent in random DNA sequences and seem to be more frequent in repeat-rich genomes (Supplemental Fig. S4) strongly suggests that repetitive DNA is the culprit, and indeed we found transposable elements and other repeats when we checked some of them. These spurious alignments are undesirable, and *k-*mer validation offers a simple and effective way to nearly eliminate them: we just have to remove from the valid *k-*mer list all *k-*mers that seem to occur more than once in the genome (see Supplemental Results: we used as a cut-off 1.5-fold of the Illumina single-copy peak; 105 in the *E. coli* case). As shown in Fig. 3, this procedure causes minimal losses in sensitivity, while suppressing most of the “false-positives”. In the next sections we will always compare the performance of standard MHAP (“M”) with the two types of *k-*mer validation: masking only low-frequency *k-*mers (“L”) or masking both low-frequency and high frequency *k-*mers (“LH”). Illumina reads allow a quite precise *k-*mer classification; given enough coverage, two-copy *k-*mers (*e.g*. from a segmental duplication) can be fairly well separated from single-copy ones (Supplemental Fig. S5). For the purpose of read correction and assembly, ideally only *k-*mers that are single-copy in the genome should be used as seeds in overlap detection; as we will see below, using Illumina reads and the modified MHAP one gets close to this.

**Figure 3.**
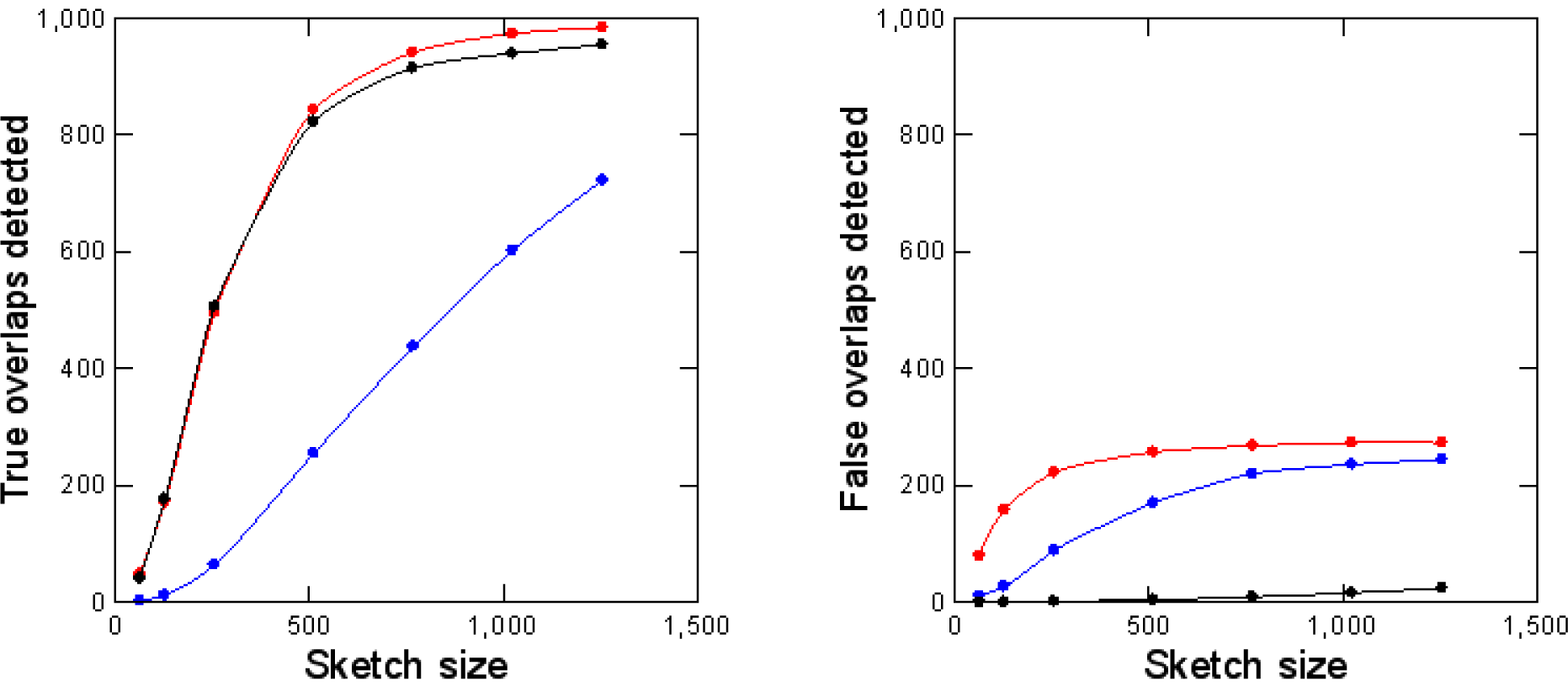
Sensitivity and specificity of read overlap detection with masking of repetitive *k-*mers. Simulated PacBio reads from *D. melanogaster* (1000 pairs of 10 kb sequences with 2kb overlaps) were subjected to standard MHAP (blue), MHAP with masking of low-frequency *k-*mers (red), or MHAP with masking of low-frequency and high-frequency *k-*mers (black). Note that masking of low and high-frequency *k-*mers cause a huge improvement in specificity (right panel) with minimal losses in sensibility (left panel). The reference list of valid *k-*mers came from Illumina reads.

### *k-*mer validation improves the correction of long-reads

We assessed the performance of read error correction by counting for each read the number of correct *k-*mers among the total *k-*mers (Supplemental Methods). Uncorrected PacBio reads from different organisms contain between 15% to 38% correct *k-*mers (Supplemental Table S1). Repeat-rich genomes have a higher proportion of correct *k-*mers possibly because errors in repetitive sequences have a rather high chance of generating a valid *k-*mer that occurs in a variant copy of the repeat (located elsewhere in the genome). During read correction in all assembly pipelines, the raw reads were aligned, regions with poor alignment were trimmed, and discrepant bases were deemed as sequencing errors and were corrected by a consensus algorithm (Chin et al. 2013; Berlin et al. 2015). Looking first at the sequencing errors (Table 1; columns 4, 7 and 10), the standard MHAP overlapper (coupled with the default falconsense correction algorithm) brings the reads from 15% -38% to 93.9% correct *k-*mers (range across different organisms: 92%-97%), and *k-*mer validation further improves this to 94.7% (L masking) and 94.8% (LH masking). Second, there are also gains in the total amount of sequence recovered (Table 1, columns 2, 5 and 8), presumably due to improved alignment and reduction of unnecessary trimming. The combined effect of these two factors is that reads corrected with LH masking have on average 221 additional correct *k-*mers (*i.e*., 15,613 minus 15,392), when compared to the standard MHAP. So *k-*mer validation indeed improves the correction of long-reads, in both trimming and error correction. The effect differs between organisms, which is expected, since it will depend on the quality of PacBio and Illumina sequencing, and on the specificities of each genome (*e.g*., amount and composition of repetitive DNA). In particular, the smallest improvement occurred in *Arabidopsis*, possibly because it has the worst Illumina dataset (Supplemental Table S2; Supplemental Fig. S1). It is interesting to also note that most of the improvement in error correction seems to be due to masking of low-frequency *k-*mers (L-masking); LH-masking (*i.e*., simultaneous masking of low-frequency and high-frequency *k-*mers) adds little in most genomes. We will return to this point later.

**Table 1.** Read error correction with different methods. All values are 95% trimmed means (to remove outliers).

| Organism | Number of reads* | Standard MHAP |  |  | L masking |  |  | LH masking |  |  |
| --- | --- | --- | --- | --- | --- | --- | --- | --- | --- | --- |
|  |  | Total *k*-mers | Correct *k*-mers | % correct | Total *k*-mers | Correct *k*-mers | % correct | Total *k*-mers | Correct *k*-mers | % correct |
| *E. coli* | 7,410 | 15,045 | 14,242 | 94.7 | 15,118 | 14,467 | 95.7 | 15,119 | 14,469 | 95.7 |
| *S. cerevisiae* | 23,455 | 11,968 | 11,016 | 92.0 | 12,122 | 11,348 | 93.6 | 12,132 | 11,377 | 93.8 |
| *C. elegans* | 128,647 | 18,701 | 17,393 | 93.0 | 18,807 | 17,566 | 93.4 | 18,806 | 17,578 | 93.5 |
| *Arabidopsis* | 185,270 | 17,429 | 16,864 | 96.8 | 17,470 | 16,973 | 97.2 | 17,468 | 16,950 | 97.0 |
| *Drosophila* | 227,987 | 18,735 | 17,461 | 93.2 | 18,826 | 17,639 | 93.7 | 18,855 | 17,705 | 93.9 |
| Grand mean | - | 16,373 | 15,392 | 93.9 | 16,466 | 15,596 | 94.7 | 16,473 | 15,613 | 94.8 |

\* Exactly the same reads were compared across the three methods.

During read correction (and assembly) we always used Illumina-derived list of valid *k-*mers, but in *E. coli* and *C. elegans* (which have completely finished genomes) we also tested the genome-derived list of valid *k-*mers to guide the read alignment. The effect in read correction is negligible (Supplemental Table S3) indicating, as seen in the previous section (Supplemental Fig. S3), that Illumina-derived lists are excellent proxies for the real *k-*mer lists.

Finally, the effect of *k*-mer validation looks small (*e.g*., 221 additional *k*-mers in 15,392, or 1.4%), but we should note that these are average values. Most assembly breaks occur at repetitive regions, and as we will see below (section Assembly of a “model genome”) at these difficult regions *k*-mer validation has a strong effect on read correction.

### *k-*mer validation results in more contiguous assemblies

We assembled the five genomes with the three assembly methods (standard MHAP, L-masking, and LH-masking), and used the Quast package (Gurevich et al. 2013) to compare them for metrics such as contiguity (NG50) and misassembly frequency (Supplemental Methods). When tested with the simple genomes of *E. coli* (4.64 Mbp) and yeast (12.1 Mbp), all three assembly methods yield similar results (Supplemental Table S4). In *E. coli*, all three approaches yield one contig spanning the complete genome, with high identity to it. In yeast, the NG50 from MHAP and LH assemblies are the same (818 kb), whereas L-masking yield a bit smaller value (751 kb). The yeast PacBio data came from W303 strain, for which there is no available finished sequence for comparison; however, the NG50 of the three assemblies approached the NG50 of the finished reference yeast strain (924 kb), so it seems that they are close to completeness. Hence both *E. coli* and yeast provide a nice demonstration of the power of long-reads which, however, leaves little room for comparison among assembly methods. However, the difference between the three assembly methods become visible in these simple genomes when we use more challenging conditions such as low coverage data or small sketch size: in both cases *k-*mer validation leads to huge improvements in assembly contiguity (Fig. 4A).

**Figure 4.**
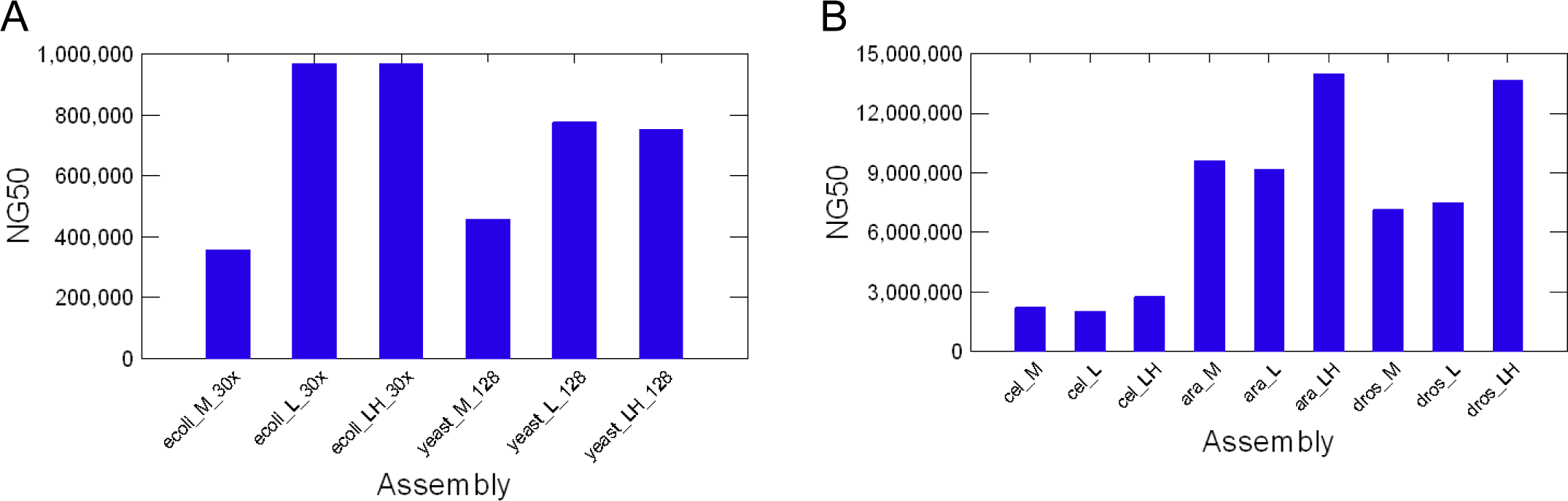
Contiguity of assemblies produced with different methods. “M”, standard MHAP; “L”, MHAP with low-frequency *k-*mer masking; “LH”, MHAP with low and high frequency *k-*mer masking. Panel A: assembly of simple genomes under the challenging conditions of low coverage (*E. coli*; coverage reduced from 94x to 30x), or small sketch size (yeast; MHAP sketch size reduced from 512 to 128). Panel B: assembly of three complex genomes (*C. elegans, A. thaliana*, and *D. melanogaster*).

When we tested the *k-*mer validation procedure with three complex genomes (*C. elegans, A. thaliana*, and *D. melanogaster*), we found that in all three cases it produced significantly more contiguous assemblies: in *C. elegans* the NG50 rose from 2,221 kb to 2,764 kb, in *Arabidopsis* from 9,588 kb to 13,994 kb, and in *Drosophila* from 7,156 kb to 13,655 (all values MHAP vs. LH-masking, Fig. 4B; Table 2). The improvement in contiguity is also seen in the largest contig size (Table 2). Statistics such as NG50 focus only on the largest contigs (*e.g*., in *Drosophila* only the 4 or 5 largest, all euchromatic), but the NGx plots indicate contiguity improvements across all size ranges (Supplemental Fig. S6). Aggressive assembly algorithms can spuriously increase statistics such as NG50 at the expense of increasing misassemblies; this was not the case of *k-*mer-validation, which actually in most cases yield smaller numbers of misassemblies, mismatches, and indels, when compared to the standard MHAP (Table 2). Although we have not tested even more complex genomes such as mammals and large plant genomes, it is very likely that *k-*mer validation will lead to improved assemblies in these cases as well.

**Table 2.**
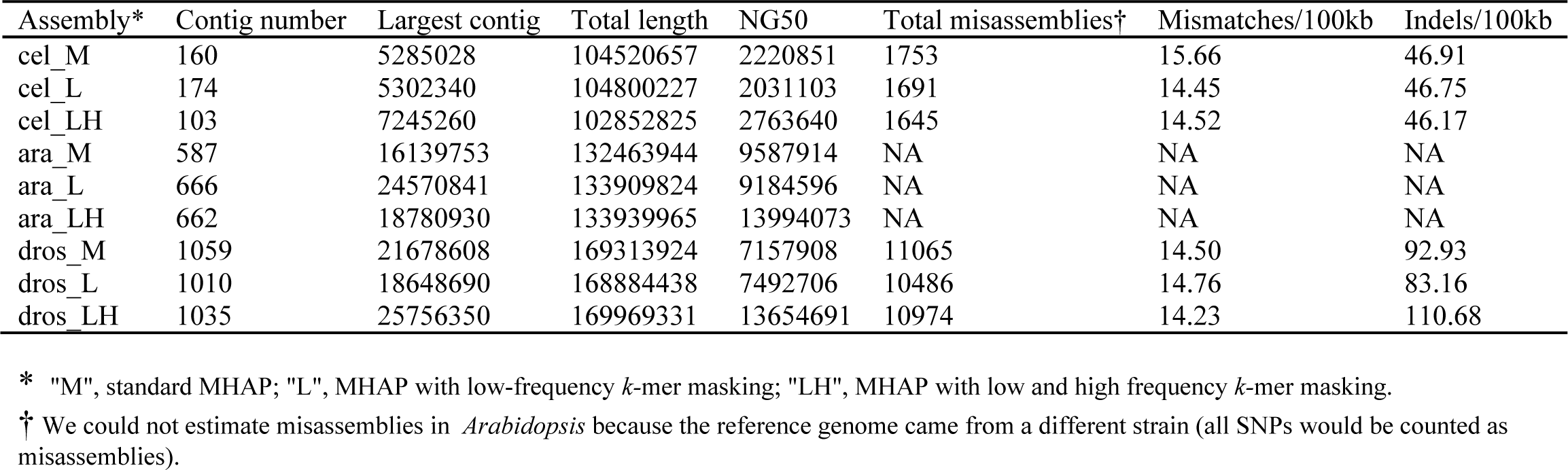
Assembly quality assessment. Note that *k-*mer validation (L and specially LH) increases the contiguity statistics (NG50, largest contig), while slightly decreasing the assembly errors (last three columns).

It is worth mentioning here two final points. First, improvements in assembly caused by *k-*mer validation are the same when we use the Illumina or the genome derived list of valid *k-*mers (Supplemental Table S5). This shows that in terms of assembly Illumina-derived lists are excellent proxies for the real *k-*mer lists, as seen before for overlap detection (Supplemental Fig. S3) and read correction (Supplemental Table S3). Second, overlap detection (Fig 3A; Supplemental Fig. S4) and read error correction (Table 1) are essentially the same with L- and LH-masking, but most or all the assembly contiguity gains in complex genomes occur with LH-masking (Fig. 4B). Hence it seems that there are additional benefits of LH-masking not fully covered by our metrics of overlap detection and read error correction. We will address this point in the next section.

### Assembly of a “model genome”

The causal events that lead to the improvements in read correction (Table 1) and in assembly (Fig. 4) are scattered in too many regions of the genome, and may too complex to allow a detailed study (*e.g*., how exactly does LH-masking improve contiguity?) In order to better understand them, we isolated a small and difficult region and used it as a model: a 44 kb segmental duplication (98% identity between the two copies), which is part of a much larger segmental duplication located in the 10q11 region of the human genome. The finished sequence of both copies was obtained by painstaking BAC cloning and sequencing (Chaisson et al. 2015). As detailed in Supplemental Methods, we used the finished sequence to simulate PacBio reads from both copies of the 44 kb segmental duplication, along with ~300 kb of flanking sequence; we used simulated reads because we want to know which segmental duplication copy they came from. We then assembled the reads with the three methods (standard MHAP, L-masking and LH-masking). In the case of L and LH-masking, we obtained the valid *k-*mer lists from the finished sequence. The perfect assembly of this “model genome” should yield two contigs (“left” and “right”), each representing one copy of the segmental duplication and the correct flanking sequences. Standard MHAP (“M”) assembly resulted in 11 contigs, L-masking in three contigs, and LH-masking yield the expected two contigs (Supplemental Table S6). The majority of the assembly breaks in the M and L assemblies were within or close to the segmental duplication region (not shown), and particularly in the M assembly there is a large amount of sequence duplication (19%), caused by partially overlapping contigs in this region (Supplemental Table S6).

Since the three assemblies differ only in the initial alignment of the uncorrected reads, all assembly differences must ultimately trace to it. When we investigated the read alignment, we found that both the standard MHAP and MHAP with L-masking fail to sort the two copies of the segmental duplication in most cases, (*i.e*., in most reads ~ 50% of the detected overlaps are between reads from different copies; Supplemental Fig. S7), whereas with LH-masking 92% of the detected overlaps are correct.

The next step in the assembly pipeline is the read correction by a consensus algorithm, using the overlaps obtained as above. Since we know the origin of reads, we can score for each site of each corrected read if it has the right base, a wrong base, or a gap. We should distinguish three types of sites here: (i) outside the segmental duplication (NSD sites); (ii) within the segmental duplication at positions that are variable between the two copies (SFV sites, for “sequence family variant”; (Dennis et al. 2012; Hughes and Rozen 2012); and (iii) within the segmental duplication at positions that are conserved between the two copies (SDC sites). Note that at SFV sites there will be conflicting sequence information in the overlaps produced by standard MHAP and by L-masking (but not by LH-masking), because as seen above these two methods mix almost indistinctly reads from the two copies of the segmental duplication. At the NSD and SDC sites, on the other hand, there is no such conflicting information, because at these sites either the two contigs do not align at all (NSD), or have the same sequence (SDC). As shown in Fig. 5, the three methods work equally well for NSD and SDC sites: in the corrected reads 98% of the bases at these sites are right. However, at SFV sites there is a huge difference: whereas with LH-masking read correction still works very well (97% right bases), with the standard MHAP and L methods only ~80% of the bases are right; in most cases the SFV site is deleted (substituted by a gap). This 80% value is the average for the whole segmental duplication; sites closer to the border actually have almost perfect correction, whereas those in the middle of segmental duplication can get below 50% correct bases (Supplemental Fig. S8). This heterogeneity in error correction makes sense: close to the border of the segmental duplication, the flanking sequence ensures the correct read overlap (and proper read correction). In the same vein, the SFV sites around a 1.5 kb indel in the middle of the segmental duplication were “protected” from miscorrection (Supplemental Fig. S8). The above results employed *falconsense* as the consensus algorithm; the more precise (and slower) *pbdagcon* yield essentially the same result (Supplemental Fig. S9), the main difference being the type of miscorrection at SFV sites: whereas *falconsense* almost always introduces a gap, *pbdagcon* do either this or introduces a wrong base. The bottom line is that in both cases the SFV information is destroyed.

**Figure 5.**
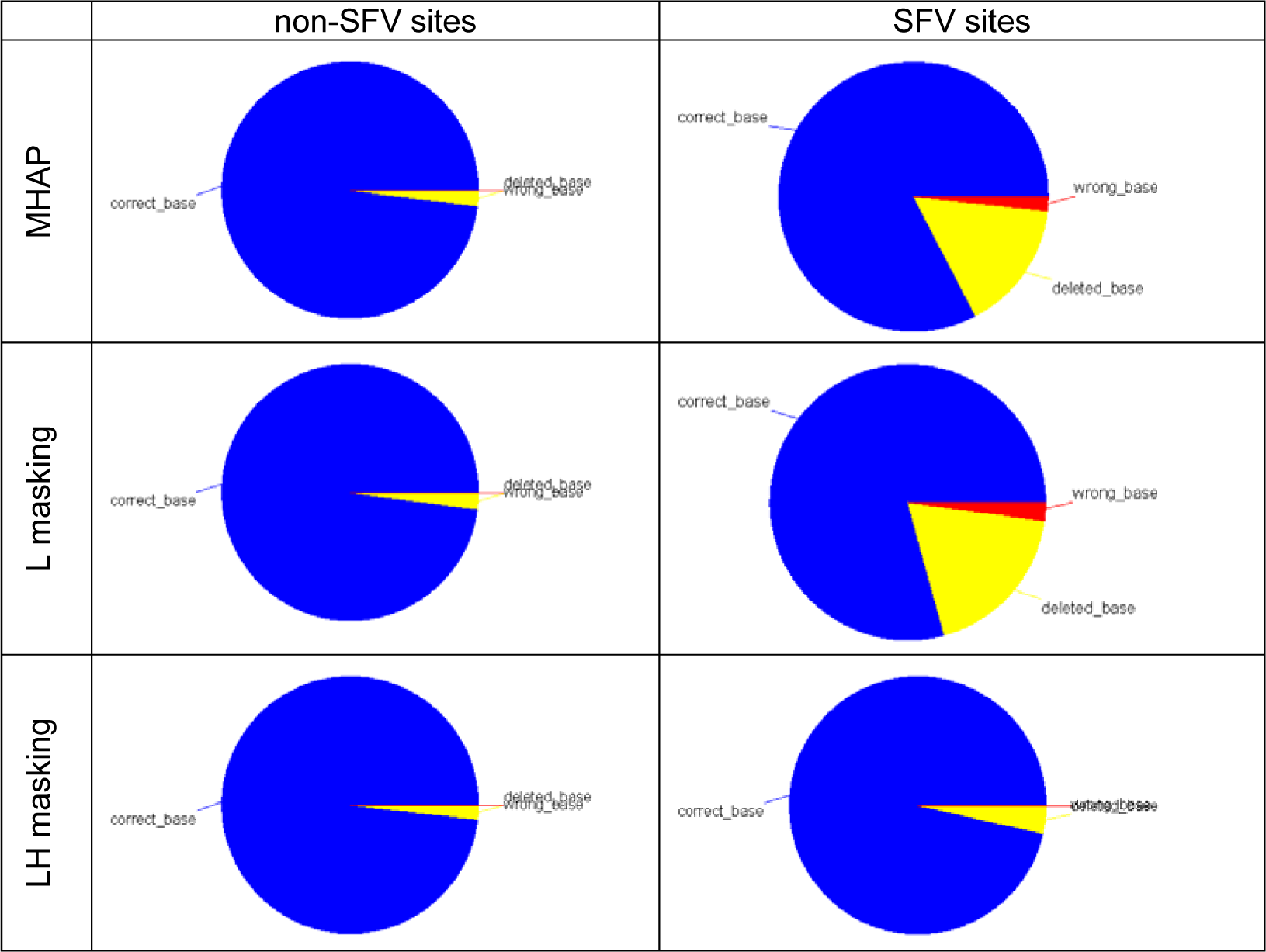
Read correction within a segmental duplication (human 10q11 region). Corrected reads were aligned with the original sequence, and each base of each read was scored as “correct” (blue), “wrong” (red), or “deleted” (yellow). “SFV sites” (for “sequence family variant”) are located within the segmental duplication, at positions where two copies are different. “Non-SFV sites” are outside the segmental duplication, or within the segmental duplication and identical between the two copies (they produce identical results and were lumped in the figure). Note that standard MHAP and MHAP with L-masking frequently fail at SFV sites, whereas LH-masking correctly handle them. Data from 450 sites of each type; reads were corrected with the default falcon-sense algorithm (see Fig. S9 for the pbdagcon correction).

So it seems that the “model genome” provided a quite complete answer for the question “how exactly does LH-masking improve contiguity? “The increase in overlap detection efficiency due to masking of error *k-*mers helps. But even more important is the stringent masking of repetitive *k-*mers (defined as all *k-*mers that are not single-copy in the genome): the different copies of a repeat can be very similar (in our example, 98% identical), and without this stringent masking the signal from SFV sites is swamped by the signal from conserved sites at the aligner step, leading first to indiscriminate overlaps (Supplemental Fig. S7), then to rampant read miscorrection at the SFV sites (Fig. 5; Supplemental Fig. S8), and finally to assembly breaks (Supplemental Table S6).

The assembly breaks are a direct consequence of the destruction of the SFV information: when a repeat is longer that the vast majority of the reads, it can only be correctly traversed by a tiling path of SFV. Ultimately, failure to correctly handle repeats during overlap detection and read correction lead to fragmentation and other assembly errors. The problems posed by repeats in genome assembly have been recognized a long time ago (Myers 1995; Phillippy et al. 2008; Nagarajan and Pop 2009; Koren et al. 2012), and long reads have a dual relationship with them: when they fully span the repeat they solve the problem, but when the repeat is longer than the reads the problem become harder because the overlap detection in principle could not be stringent. In a sense, LH-masking implements stringent overlap detection in noisy reads.

### Sampling bias in the *Drosophila* PacBio data

Given previous work that showed that PacBio sequencing solved two difficult repetitive regions of the *Drosophila* Y-chromosome (Carvalho et al. 2015; Krsticevic et al. 2015), we were surprised to find that some Y-linked single-copy genes were missing large parts (*e.g*., only 24% of the coding sequence of the *kl-5* gene was present in the original MHAP assembly; Fig. 6A; Supplemental Table S7).

**Figure 6.**
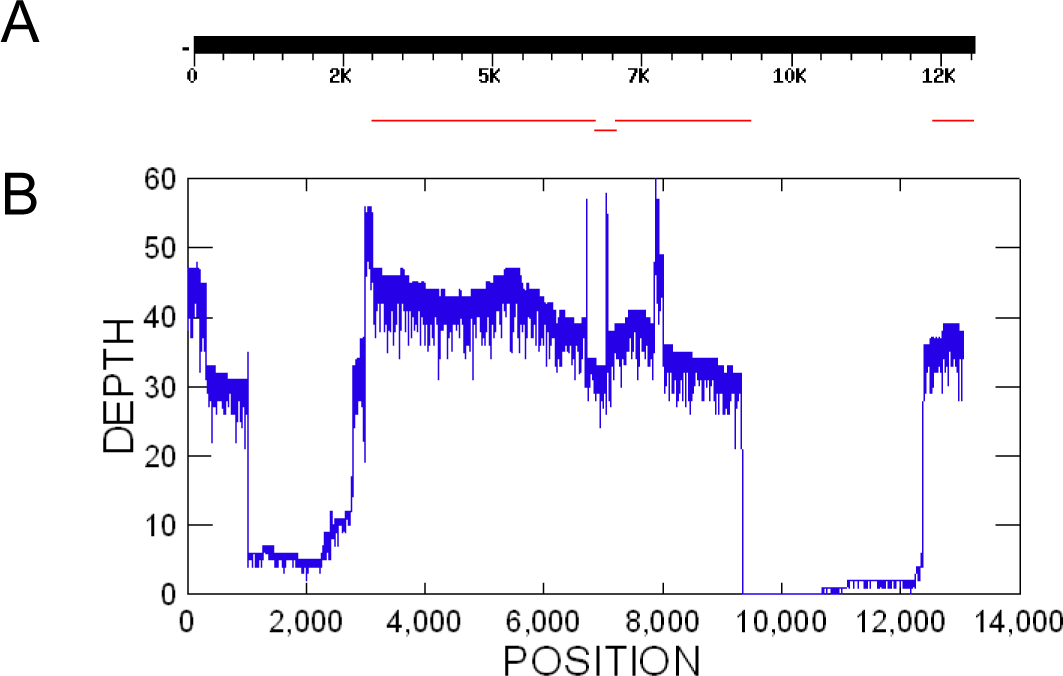
Sampling bias in the *Drosophila* PacBio data. Panel A: BlastN search using the Y-linked *kl-3* gene CDS as the query against a database of the MHAP-assembled *Drosophila* genome (Berlin et al. 2015). Panel B: Coverage of the same gene by raw PacBio reads. Note that most of assembly gaps in the *kl-3* gene actually were caused by low or absent coverage by PacBio reads. The expected read coverage is 45x. The Illumina coverage of the same region is fairly homogeneous (Fig. S10).

We initially thought that this was due to a combination of the lower coverage of the Y (~ 45x; the *Drosophila* reads came from male DNA, and hence coverage of the sex-chromosomes should be half of the autosomes), and assembly parameters optimized for the ~95x coverage of the autosomes. Two findings of the present work strongly suggest that this was not the correct explanation: (i) *k-*mer validation, and tweaking of assembly parameters, caused only marginal improvements (we tried around 40 different combinations of the parameters ovlMinLen, merThreshold, and assembleCoverage; data not shown); (ii) the coding regions of all 20 X-linked genes we looked are complete (Supplemental Table S7). When we looked at the raw PacBio reads, we found that sequencing depth was very irregular in several Y-linked genes, reaching nearly zero in large parts of *kl-3, kl-5* and another genes, whereas the sequencing depth of X-linked genes is fairly constant and centered around 45x, as expected (Supplemental Fig. S10).

This finding is important and may have general significance because it violates one of the conditions of Gene Myers “140 char theorem” (“*sampling is Poisson* “); such violations of random sampling may be an obstacle to perfect assemblies using PacBio technology.

The strong sequencing bias described above is surprising, given the success of PacBio data in assembling AT-rich or GC-rich genomes (Shin et al. 2013; Paredes et al. 2015) and previous reports of fairly uniform coverage across genomes (Ross et al. 2013). We believe that this bias is related to the peculiar organization of some *Drosophila* Y-linked genes, which have Mbp-sized introns composed of simple satellite DNA (*e.g*. (AT)_n_; the location of these satellite blocks is not precisely known; (Bonaccorsi and Lohe 1991; Kurek et al. 2000; Reugels et al. 2000)). This will not cause problems in the assembly of exons with Sanger or short-read technologies, because the DNA is sheared in short pieces before sequencing. Indeed, the Illumina coverage is fairly constant across all Y-linked genes (except for occasional exon duplications; Supplemental Fig. S10). However, DNA used for PacBio sequencing has high molecular weight, in the ~100 kb range when extracted and then sheared to ~20kb-40 kb; this means that some Y-linked exons will always be surrounded by a large chunk of simple repeats. Indeed, when we looked at the exon with the lowest coverage in the *kl-5* gene, we found that it is surrounded by at least ~10 kb of nearly pure (AT)_n_ sequence on one side, and a very AT-rich sequence on the other side (Supplemental Fig. S11).

How might had these repeats adversely affected PacBio sequencing? A benign hypothesis would be at the sample preparation step: (Kim et al. 2014) reported the use of cesium chloride centrifugation for the *Drosophila* sample, which may had selected against AT-rich regions such as exons flanked by massive AT-rich satellite blocks (they will have a smaller buoyant density). A more worrisome possibility is that PacBio sequencing has some intrinsic, strong bias (*e.g*., against regions with very strong AT-bias). One way to solve the question would be to sequence again *D. melanogaster*, without the use of cesium chloride centrifugation for sample preparation. It is ironic that we failed to improve the assembly of single-copy Y-linked genes from *Drosophila*, since this was the original motivation of the present work.

## DISCUSSION

Single molecule sequencing is revolutionizing genome assembly: the long reads can yield Mbp-sized contigs that span complete chromosomes (or nearly so) of prokaryotes and simple eukaryotes, and the euchromatic parts of more complex genomes such as *Drosophila* (Berlin et al. 2015; Koren and Phillippy 2015). Their major limitation is the low accuracy. Specifically, the high error rate generates a huge number of *k-*mers that are not present in the original genome, and the aligners (*e.g*., MHAP) must sift through them in order to find shared, real *k-*mers that indicate true read overlaps. These problems currently are addressed by sequencing at high depth (ideally 100x), aligning the reads with improved, fast software (Myers 2014; Berlin et al. 2015), and implementing a consensus algorithm to correct the reads prior to normal assembly (Chin et al. 2013). Whereas these procedures in principle are straightforward, the computational cost is high, and can be nearly prohibitive for large genomes (*e.g*., mammals). A less appreciated problem is the risk of miscorrection of the reads from repetitive regions: as the initial alignment must be loose in order to detect real overlaps among the noisy reads, reads from paralogous regions (*e.g*., different copies of tandem rDNA genes, long transposons, or segmental duplications) will easily be lumped together (Supplemental Fig. S7); once this happens, the error correction algorithm miscorrect the reads at the “sequence family variant” sites (Fig. 5), which in later assembly steps tend to cause assembly breaks.

In this paper we propose a simple and inexpensive procedure that addresses both problems: to enforce that only correct, single-copy *k-*mers are used as seeds for the read alignment. The enforcing of “correct *k-*mers” solves the “needle in a haystack problem”, by making the aligner ignore the error *k-*mers, which are the vast majority. This by itself dramatically increase the sensibility in overlap detection of the MHAP aligner (Fig. 2). The enforcing of *k-*mers that are single-copy in the genome increases the specificity in read overlapping (Fig 3B; Supplemental Fig. S7) and essentially abolish read miscorrection at “sequence family variant” sites (Fig. 5). This procedure requires a list of all *k-*mers from the genome. Whereas a perfect list can only be obtained from a completely finished genome (*i.e*., when a new assembly is nonsensical), we showed that *k-*mers from Illumina reads provide an excellent approximation to it. In contrast to the direct correction of PacBio reads with Illumina reads (“hybrid assemblies”), we used Illumina reads only as a source of the list of correct single-copy *k-*mers. This list is used to inform the aligner which *k-*mers should be ignored, thus guiding the alignment of PacBio reads for their self-correction; all sequence information came from the PacBio reads themselves. Its use significantly improves overlap detection (Fig. 2), the accuracy of read correction (Table 1; Fig. 5; Supplemental Fig. S8), and the contiguity and accuracy of genome assembly (Fig. 4; Table 2). Gains in contiguity as measured by NG50 ranged from 24% (in *C. elegans*) to 91% (*i.e*., almost doubled, in *D. melanogaster*). We believe that these gains justify by themselves the use *k-*mer validation, but larger gains may be possible, as suggested below. Finaly, note that the additional cost of Illumina sequencing is negligible, or even absent, since in nearly all cases in which a PacBio dataset is available, there is a also Illumina dataset from the same strain. In cases where one needs to do the Illumina sequencing, Supplemental Fig. S1 and Supplemental Table S2 suggest that a ~70x coverage is enough for a good separation between single-copy and repetitive *k*-mers.

### Possible improvements

There are several avenues that may lead to further improvements in assembly contiguity. First, nearly all current long read assemblers employ two rounds of read alignment and correction (the first deals with the raw reads, and the second with the correct reads), and we modified only the aligner of the first round (the MHAP program). It is likely that the second round of read alignment and correction, which uses a different program and is aimed to deal with Sanger-like error rates, would also benefit from the stringent use of single-copy *k-*mers as seeds for overlap detection.

Second, it will be interesting to test the *k-*mer validation procedure in the other long-read aligners (DALIGNER and BLASR) because MHAP is a “probabilistic” aligner that randomly samples a subset of *k-*mers, whereas both DALINGER and BLASR deal with all *k-*mers. It is likely that *k-*mer validation will increase the speed (and possibly the stability) of these aligners.

Third, the read correction algorithms (*e.g*., *falconsense* and *pbdagcon*), may be modified to avoid the introduction of non-valid *k-*mers while correcting the reads.

Fourth, there is the issue of the Illumina-derived *k-*mer list. As detailed in the Supplemental Results, we did a limited exploration of the low frequency cut-off (which should remove error *k-*mers) and high frequency cut-off (which should spare only single-copy *k-*mers), and hence a more throughout trial-and-error may be beneficial.

Fifth, nearly uniform coverage is considered a major advantage of PacBio sequencing, and hence the strong bias we found in the *Drosophila* data warrants further investigations. It will also be interesting to look at Oxford Nanopore datasets for complex genomes, which as far as we know have not yet been produced.

Finally, very recently the PBcR/Celera Assembler pipeline has been upgraded to a new pipeline called Canu (http://canu.readthedocs.org/). It will be very interesting to implement the *k-*mer validation procedure on it.

These developments are beyond the scope of the present paper; we are pursuing some of them now.

### How far can we go?

Assembly quality is a function of coverage, error rate, and read length (Phillippy et al. 2008; Nagarajan and Pop 2009; Myers 2016). Second generation technology (*e.g*. Illumina) provided a good solution for this equation when fragmentation (and correct repeat reconstruction) is not a concern, *e.g*., for sequencing genes or to identify SNP variants by comparison to a reference genome (The 1000 Genomes Project Consortium 2010). Long read sequencing provided a different solution: it yields unfragmented, nearly finished assemblies of regions with moderate repeat content, such as prokaryotic genomes and (to a large extent) the euchromatic portion of complex eukaryotic genomes, at a higher cost. However, sequencing costs of new technologies tend to drop quickly, and the maturation of other long read technologies (*e.g*., Oxford Nanopore) brings the promise of further cost reductions. Hence, the major challenge that remains is how to correctly assemble repetitive DNA, which currently cause large assembly gaps (*e.g*., the histone and rDNA clusters of *Drosophila*, nearly all centromeres), massive fragmentation in the heterochromatin, and scattered breaks in the euchromatin (*e.g*., humans segmental duplications). As (Myers 2016) stressed, this is an open question: “… work on the assembly problem has failed to really address the issue of how to resolve repetitive sequences except in fairly superficial ways.” In a sense, technology development is pushing forward what a is repeat in assembly terms (“*reads long enough 2 solve repeats* “): retrotransposons (~7kb long) are a major obstacle for contig building with Sanger and Illumina sequencing, and are almost harmless to PacBio. So brute force, in the form of very long reads (say, average length in the 100 kb range), would solve the majority of the currently intractable regions mentioned above: once the “golden threshold” of reads-longer-than-repeats is crossed, genome assembly became much simpler (see Fig. 1 in (Koren and Phillippy 2015)).

But “perfect assembly” is possible even when reads are not long enough to cross a repeat, as SFVs may provide a unique tiling path across it. For example, no read used in the assembly of our “model genome” spans the 44 kb segmental duplication, and yet we could assemble it in a essentially perfect form. As the results from our “model genome” show, the key is first to preserve the SFV sites by not miscorrecting them, and second, to effectively use their information, by not swamping the overlap detection with the flood of repetitive (*i.e*., non single-copy) *k-*mers. The *k-*mer validation procedure we presented here seems to be an effective implementation of these two features. Ultimately the ability to cross a repeat longer than the read length will depend on the number of SFVs per read. A tiling path requires an absolute minimum of two SFVs per read, and our model genome data had roughly 141 SFVs per read (450 SFV sites in a 44kb segmental duplication; average length of corrected reads: 13767 bp). It remains to be seen which read length will provide enough SFVs to cross large regions such as the histone or rDNA clusters in *Drosophila* (500 kb and 2 Mbp, respectively) which currently are inaccessible (both are severely fragmented even in our best *Drosophila* assembly). In the same vein, two gaps in (Chaisson et al. 2015) could not be closed with BAC-based assembly, which amount to reads in the ~100 kb range, with very low error rate. Another limit, admittedly secondary, is the assembly of simple repeats such as the intronic (AT)_n_ blocks of *Drosophila* Y-inked genes, because the repeat periodicity (2-10 bp) overlaps with the error frequency of the uncorrected long reads. Finally, *k*-mer validation (with LH-masking) is useful even when repeats are smaller than the read length, for it protects the reads from miscorrection at repeats and hence reduce assembly errors in these regions.

As sequencing technology and assembly software move forward, the question posed by the title of this section keeps returning (Weber and Myers 1997; Carvalho et al. 2003; Koren and Phillippy 2015; Myers 2016). But as clearly stated by Myers (https://dazzlerblog.wordpress.com/2014/05/15/on-perfect-assembly/) and (Koren and Phillippy 2015), perfect assemblies are on the verge of becoming reality, and we may now be close to the final answer.

## METHODS

### Sequence reads

The sources of PacBio and Illumina reads for all five organisms are shown in Supplemental Table S8. All PacBio reads came from (Kim et al. 2014) and PacBio DevNet (http://www.pacb.com/); we downloaded them from Amazon S3 repositories (listed in the Supplementary Information of (Kim et al. 2014), or from the Amazon Elastic Block Storage (EBS) snapshot described in (Berlin et al. 2015). These data have also been deposited at NCBI Short Read Archive (except for *C. elegans*), but the reads there are unfiltered (Kim et al. 2014). The sources of Illumina reads follows: *E. coli*, Illumina BaseSpace (https://basespace.illumina.com); *S. cerevisae*, Saccharomyces Genome Database (http://www.yeastgenome.org/); *A. thaliana*, (Cao et al. 2011; Gan et al. 2011); *C. elegans*, (van Schendel et al. 2015); *D. melanogaster*, (Gutzwiller et al. 2015).

### Implementation of *k-*mer validation

We implemented the *k-*mer validation in the MHAP overlapper as follows. The standard MHAP algorithm converts each *k-*mer to a number (using a hash function), and saves from each read only the lowest value (called “min-mer”). The process is repeated, say, 500 times with different hash functions to generate a “sketch” of size 500, which is stored in the memory; overlapping reads were detected because their sketches share min-mers above a user-specified cut-off (see (Berlin et al. 2015) for details). We implemented the *k-*mer validation by adding a simple step in the MHAP code: if the read *k-*mer is present in the valid *k-*mer list, it is converted to a number as described above. If it is not there (and hence probably is a an error *k-*mer), it is converted to a very large number (technically, to Long.MAX_VALUE), effectively forcing the program to ignore it. The list of valid *k-*mers was previously obtained from Illumina reads with the *jellyfish* program (Marçais and Kingsford 2011), saved as a text file, and read by the modified MHAP code, before reading the PacBio reads (see the README.txt file in http://tinyurl.com/modified-MHAP). Note that these procedures implement a “positive selection list”, whereas most aligners and overlappers allow for a list of undesirable *k-*mers (either supplied by the user or produced by the program itself) which is used to remove highly repetitive *k-*mers, in order to reduce the computational load (*e.g*., the “*filter-threshold*” parameter in MHAP). Furthermore, their identification of repetitive *k-*mers is much less precise because *k-*mer counts are obtained from the raw PacBio reads. Illumina reads allow a much finer *k-*mer classification; given enough coverage, even two-copy *k-*mers (*e.g*. from a segmental duplication) can be fairly well separated from single-copy ones (Supplemental Fig. S5). The frequency cut-off values used to build the valid *k-*mer lists is presented in Supplemental Table S2 and Supplemental Fig. S1, and further discussed in the Supplemental Results.

The modified MHAP (source and compiled jar file) and PBcR script are available at http://tinyurl.com/modified-MHAP. The same link provides a README.txt file, with instructions on how to install and run the modified files. When run without a valid *k-*mer list the modified MHAP produces an output that is identical to the original MHAP code.

The same list of valid *k*-mers mentioned above was used to sort “correct” and “error” *k*-mers in reads. For example, in Fig.1 we used a custom script that loaded the list in the memory (as an associative array) and used it to classify each read *k*-mer as correct (match) or error (not match).

### Genome assemblies

All assemblies were performed in two Linux servers with 24 cores and 64 Gb or 144 Gb of RAM, with the Celera Assembler version 8.3 (PBcR pipeline). Unless otherwise noted, default PBcR parameters were used for all assemblies, including the *falconsense* read correction algorithm. Our main purpose was to compare the Standard MHAP overlapper with the modified version (*i.e*., with *k-*mer validation) and to save time we opted to not polish the assemblies with Quiver (Chin et al. 2013).

## DATA ACCESS

The software developed in this work is available at http://tinyurl.com/modified-MHAP. The resulting genome assemblies of *C. elegans, Arabidopsis* and *Drosophila* are available at http://tinyurl.com/assembled-genomes.

## ACKNOWLEDGMENTS

We thank R Hoskins and S. Celniker for first calling our attention to PacBio sequencing, Carlos Martins and Paulo Abdon for java programming, and R. Hoskins, B. Lemos, L. Koerich, M. Vibranovski, S. Koren, and our lab members for valuable suggestions in the manuscript and help. We thank Reinaldo Brito (UFSCAR), Regis Corrêa and Gilberto Sachetto (UFRJ) for granting us access to their servers. This work was supported by Conselho Nacional de Desenvolvimento Científico e Tecnológico-CNPq and Fundação de Amparo à Pesquisa do Estado do Rio de Janeiro-FAPERJ.

## AUTHOR CONTRIBUTIONS

A.B.C. conceived the work, performed the research, analyzed the data and wrote the manuscript; E.G.D. and G.N. performed the research and analyzed the data.

## REFERENCES

Berlin K, Koren S, Chin CS, Drake JP, Landolin JM, Phillippy AM. 2015. Assembling large genomes with single-molecule sequencing and locality-sensitive hashing. Nature biotechnology 33(6): 623–630.

Blattner FR, Plunkett G, 3rd, Bloch CA, Perna NT, Burland V, Riley M, Collado-Vides J, Glasner JD, Rode CK, Mayhew GF et al. 1997. The complete genome sequence of Escherichia coli K-12. Science 277(5331): 1453–1462.

Bonaccorsi S, Lohe A. 1991. Fine mapping of satellite DNA-sequences along the Ychromosome of Drosophila melanogaster - Relationships between satellite sequences and fertility factors. Genetics 129(1): 177–189.

Cao J, Schneeberger K, Ossowski S, Gunther T, Bender S, Fitz J, Koenig D, Lanz C, Stegle O, Lippert C et al. 2011. Whole-genome sequencing of multiple Arabidopsis thaliana populations. Nat Genet 43(10): 956–963.

Carvalho AB, Vibranovski MD, Carlson JW, Celniker SE, Hoskins RA, Rubin GM, Sutton G, Adams M, Myers EW, Clark AG. 2003. Y chromosome and other heterochromatic sequences of the Drosophila melanogaster genome; how far can we go? Genetica 117: 227–237.

Carvalho AB, Vicoso B, Russo CA, Swenor B, Clark AG. 2015. Birth of a new gene on the Y chromosome of Drosophila melanogaster. Proc Natl Acad Sci U S A 112(40): 12450–12455.

Chaisson MJ, Huddleston J, Dennis MY, Sudmant PH, Malig M, Hormozdiari F, Antonacci F, Surti U, Sandstrom R, Boitano M et al. 2015. Resolving the complexity of the human genome using single-molecule sequencing. Nature 517(7536): 608–611.

Chaisson MJ, Tesler G. 2012. Mapping single molecule sequencing reads using basic local alignment with successive refinement (BLASR): application and theory. BMC bioinformatics 13: 238.

Chin CS, Alexander DH, Marks P, Klammer AA, Drake J, Heiner C, Clum A, Copeland A, Huddleston J, Eichler EE et al. 2013. Nonhybrid, finished microbial genome assemblies from long-read SMRT sequencing data. Nat Methods 10(6): 563–569.

Dennis MY, Nuttle X, Sudmant PH, Antonacci F, Graves TA, Nefedov M, Rosenfeld JA, Sajjadian S, Malig M, Kotkiewicz H et al. 2012. Evolution of human-specific neural SRGAP2 genes by incomplete segmental duplication. Cell 149(4): 912–922.

Drosophila_Community_Resources_Committee. 2001. Drosophila White Paper 2001.

Gan X, Stegle O, Behr J, Steffen JG, Drewe P, Hildebrand KL, Lyngsoe R, Schultheiss SJ, Osborne EJ, Sreedharan VT et al. 2011. Multiple reference genomes and transcriptomes for Arabidopsis thaliana. Nature 477(7365): 419–423.

Goodwin S, Gurtowski J, Ethe-Sayers S, Deshpande P, Schatz MC, McCombie WR. 2015. Oxford Nanopore sequencing, hybrid error correction, and de novo assembly of a eukaryotic genome. Genome Res 25(11): 1750–1756.

Gurevich A, Saveliev V, Vyahhi N, Tesler G. 2013. QUAST: quality assessment tool for genome assemblies. Bioinformatics 29(8): 1072–1075.

Gutzwiller F, Carmo CR, Miller DE, Rice DW, Newton IL, Hawley RS, Teixeira L, Bergman CM. 2015. Dynamics of Wolbachia pipientis Gene Expression Across the Drosophila melanogaster Life Cycle. G3 5(12): 2843–2856.

Hughes JF, Rozen S. 2012. Genomics and genetics of human and primate Y chromosomes. Annu Rev Genomics Hum Genet 13: 83–108.

Kim KE, Peluso P, Babayan P, Yeadon PJ, Yu C, Fisher WW, Chin C-S, Rapicavoli NA, Rank DR, Li J. 2014. Long-read, whole-genome shotgun sequence data for five model organisms. Scientific Data 1: 140045.

Koren S, Harhay GP, Smith TP, Bono JL, Harhay DM, McVey SD, Radune D, Bergman NH, Phillippy AM. 2013. Reducing assembly complexity of microbial genomes with single-molecule sequencing. Genome Biol 14(9): R101.

Koren S, Phillippy AM. 2015. One chromosome, one contig: complete microbial genomes from long-read sequencing and assembly. Current opinion in microbiology 23: 110–120.

Koren S, Schatz MC, Walenz BP, Martin J, Howard JT, Ganapathy G, Wang Z, Rasko DA, McCombie WR, Jarvis ED et al. 2012. Hybrid error correction and de novo assembly of single-molecule sequencing reads. Nature biotechnology 30(7): 693–700.

Krsticevic FJ, Schrago CG, Carvalho AB. 2015. Long-read single molecule sequencing to resolve tandem gene copies: The Mst77Y region on the Drosophila melanogaster Y chromosome. G3 5(6): 1145–1150.

Kurek R, Reugels AM, Lammermann U, Bunemann H. 2000. Molecular aspects of intron evolution in dynein encoding mega- genes on the heterochromatic Y chromosome of Drosophila sp. Genetica 109(1-2): 113–123.

Li H. 2016. Minimap and miniasm: fast mapping and de novo assembly for noisy long sequences. Bioinformatics.

Loman NJ, Quick J, Simpson JT. 2015. A complete bacterial genome assembled de novo using only nanopore sequencing data. Nat Meth 12(8): 733–735.

Marçais G, Kingsford C. 2011. A fast, lock-free approach for efficient parallel counting of occurrences of k-mers. Bioinformatics 27(6): 764–770.

McCoy RC, Taylor RW, Blauwkamp TA, Kelley JL, Kertesz M, Pushkarev D, Petrov DA, Fiston-Lavier AS. 2014. Illumina TruSeq synthetic long-reads empower de novo assembly and resolve complex, highly-repetitive transposable elements. PLoS One 9(9): e106689.

Myers EW. 1995. Toward simplifying and accurately formulating fragment assembly. Journal of computational biology: a journal of computational molecular cell biology 2: 275–290.

Myers G. 2014. Efficient local alignment discovery amongst noisy long reads. In Algorithms in Bioinformatics, pp. 52–67. Springer.

Myers G. 2016. A history of DNA sequence assembly. Information Technology. in press

Nagarajan N, Pop M. 2009. Parametric complexity of sequence assembly: theory and applications to next generation sequencing. Journal of computational biology: a journal of computational molecular cell biology 16(7): 897–908.

Nagarajan N, Pop M. 2013. Sequence assembly demystified. Nat Rev Genet 14(3): 157–167.

Paredes JC, Herren JK, Schupfer F, Marin R, Claverol S, Kuo CH, Lemaitre B, Beven L. 2015. Genome sequence of the Drosophila melanogaster male-killing Spiroplasma strain MSRO endosymbiont. mBio 6(2).

Phillippy AM, Schatz MC, Pop M. 2008. Genome assembly forensics: finding the elusive mis-assembly. Genome Biol 9(3): R55.

Reugels AM, Kurek R, Lammermann U, Bunemann H. 2000. Mega-introns in the dynein gene DhDhc7(Y) on the heterochromatic Y chromosome give rise to the giant Threads loops in primary spermatocytes of Drosophila hydei. Genetics 154(2): 759–769.

Ross MG, Russ C, Costello M, Hollinger A, Lennon NJ, Hegarty R, Nusbaum C, Jaffe DB. 2013. Characterizing and measuring bias in sequence data. Genome Biol 14(5): R51.

Shin SC, Ahn do H, Kim SJ, Lee H, Oh TJ, Lee JE, Park H. 2013. Advantages of Single-Molecule Real-Time Sequencing in High-GC Content Genomes. PLoS One 8(7): e68824.

The 1000 Genomes Project Consortium. 2010. A map of human genome variation from population-scale sequencing. Nature 467(7319): 1061–1073.

van Schendel R, Roerink SF, Portegijs V, van den Heuvel S, Tijsterman M. 2015. Polymerase [Theta] is a key driver of genome evolution and of CRISPR/Cas9-mediated mutagenesis. Nature communications 6.

Weber JL, Myers EW. 1997. Human whole-genome shotgun sequencing. Genome Research 7(5): 401–409.

